# Chromosome-level genome assemblies of the red algae *Porphyra dioica* and *Porphyra linearis*

**DOI:** 10.64898/2026.05.14.725108

**Authors:** Jordi Morcillo, Sofie D’hondt, Agnieszka P. Lipinska, Silke Bouckenooghe, Louka Noyen, Antoine Van de Vloet, Sofie Vranken, Jessica Knoop, Frederik Leliaert, Olivier De Clerck

## Abstract

As one of the earliest-diverging multicellular eukaryotic lineages, the bladed Bangiales (Rhodophyta) possess a deep evolutionary history with a central role in the multi-billion-dollar global seaweed aquaculture industry. Although North Atlantic representatives are emerging candidates for regional mariculture, the scarcity of high-quality genomic resources for these taxa hinders both fundamental research and commercial optimization. To address this, we present the first chromosome-level genome assemblies for two native European species: *Porphyra dioica* (150.44 Mbp) and *Porphyra linearis* (95.22 Mbp). By integrating Oxford Nanopore Technologies (ONT) long-read sequencing with Hi-C proximity ligation, we generated highly contiguous nuclear genomes resolved into five chromosomes. Structural gene models were predicted through the BRAKER3 pipeline, identifying 12,548 and 10,382 protein-coding genes for *P. dioica* and *P. linearis*, respectively. Subsequent homology-based functional annotation characterized 57.4% and 59.8% of these predicted proteins. Supplemented by circularized organellar genomes, these reference genomes provide a critical framework for future research, enabling comparative studies of Atlantic-Pacific divergence and facilitating the development of selective breeding programs for sustainable European aquaculture.

## Background & Summary

Red algae (Rhodophyta) represent one of the most ancient eukaryotic lineages, possessing a fossil record that extends beyond 1.6 billion years. Precambrian representatives, including *Rafatazmia* (∼1.6 Ga), *Ramathallus* (∼1.6 Ga) and *Bangiomorpha* (∼1.0 Ga), offer critical evidence into the initial emergence of complex multicellularity^1-3^. Examination of these structural remnants reveals filamentous architectures, cell to cell junctions consistent with modern pit connections, and differentiated cells interpreted as primitive sexual reproductive structures, providing rare insight into the early evolution of cellular organization and sexual reproduction among eukaryotes^4,5^. Extant members of the Bangiales, such as the heteromorphic genera *Porphyra* C.Agardh and *Pyropia* J.Agardh, retain these ancestral traits *—* most notably during the conchocelis phase *—* alongside specialized physiological adaptations that enable them to dominate the extreme and fluctuating environments of the intertidal zone. In these habitats, they successfully cope with recurrent desiccation, acute osmotic stress, intense irradiance, and large thermal fluctuations^6-8^.

Beyond their evolutionary significance, bladed Bangiales are fundamental in global seaweed aquaculture. In East Asia, the mariculture of species like *Neopyropia yezoensis* (Ueda) L.-E.Yang & J.Brodie has matured from its traditional foundations in the late 17th century^9^ into the current sophisticated, multi-billion-dollar industry for *nori* production^10^. Conversely, the commercial exploitation of native North Atlantic species *—* such as *Porphyra dioica* J.Brodie & L.M.Irvine or *Porphyra linearis* Greville *—* remains under active refinement^11,12^. Intriguingly, recent research on European *Porphyra* species has revealed substantial biological complexity, characterized by intricate life cycles, chimeric blades, cryptic speciation, polyploidy and interspecific hybridization^13-15^. However, the reliance on a limited set of genetic markers^16^ has left fundamental gaps in our understanding of their diversity, life cycle variation and evolutionary history. Such observations underscore the urgent requirement for comprehensive genomic resources to elucidate the mechanisms governing life-history transitions, resolve ambiguous phylogenetic relationships, and provide the necessary framework for a sustainable development of the North Atlantic seaweed sector.

Despite significant advances in sequencing technologies, genomic resources for macroalgae remain disproportionately limited. The generation of reference-quality nuclear genome assemblies is frequently hampered by the challenging biochemical properties of macroalgal tissues. For instance, complex cell-wall polysaccharides — such as galactans in the Rhodophyta, alginates in the Phaeophyceae, and ulvans in the Chlorophyta — often co-purify with nucleic acids, putatively compromising the recovery of good-quality, High Molecular Weight (HMW) DNA. In brown lineages, this challenge is compounded by high concentrations of polyphenols, which can form covalent DNA adducts and critically diminish the performance of long-read sequencing platforms^17^. Moreover, the prevalence of dense epiphytic bacterial communities and high-copy organellar genomes (chloroplasts and mitochondria) can represent a significant sequencing burden, often accounting for 70–80% of total reads^18^. Even rigorous decontamination, such as antibiotic treatments or mechanical surface abrasion, often fail to fully eliminate microbial DNA, making the assembly of highly contiguous genomes particularly challenging^6-8^. As a result, chromosome-level assemblies within the Bangiales are limited thus far to two East Asian species: *Neopyropia yezoensis* (∼108 Mb, 3 chromosomes)^7^ and *Neoporphyra haitanensis* (T.J.Chang & B.F.Zheng) J.Brodie & L.-E.Yang (∼50 Mb, 5 chromosomes)^8,19^. The current benchmark for the genus *Porphyra* is a scaffold-level assembly of a North American *P. umbilicalis* isolate (∼88 Mb across 2,125 scaffolds)^6^, with no whole-genome sequences currently available for European lineages.

To address this gap, we present high-quality, chromosome-level genome assemblies of *Porphyra dioica* and *Porphyra linearis*, generated as part of the European Reference Genome Atlas (ERGA) initiative within the Biodiversity Genomics Europe (BGE) project. By integrating Oxford Nanopore Technologies (ONT) long-read sequencing with Hi-C proximity ligation, we produced two highly contiguous nuclear genome assemblies anchored to chromosomal-scale scaffolds. These are complemented by complete, circularized mitochondrion and chloroplast genomes, establishing a comprehensive genomic baseline for North Atlantic *Porphyra* species. To facilitate downstream comparative and functional studies, both the nuclear and organellar architectures were supplemented with gene models and functional annotations. A schematic overview of our sequencing, assembly, and annotation workflow is provided in Fig. 1.

**Figure 1.**
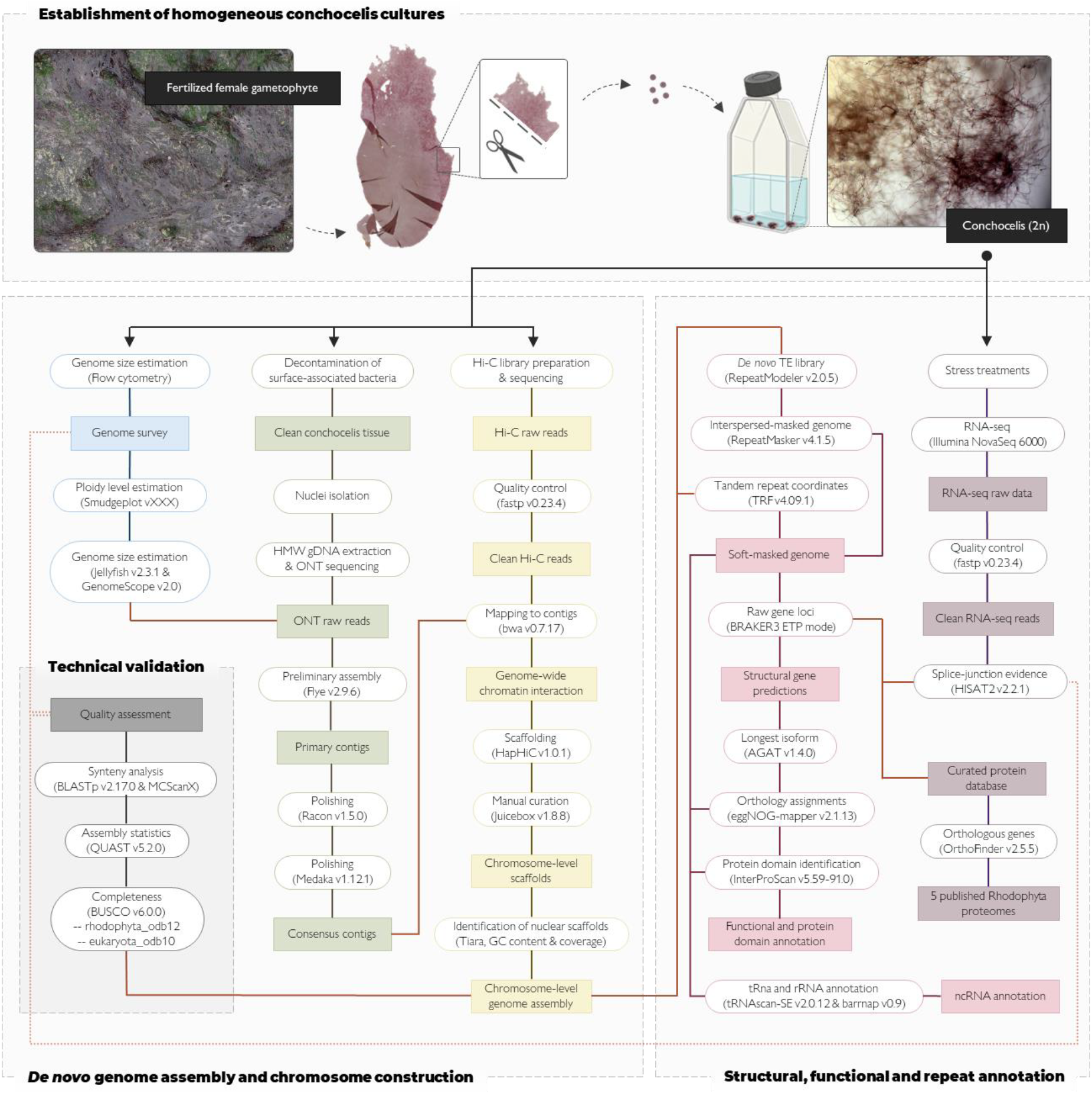
Integrated workflow for the generation, annotation, and validation of chromosome-level *Porphyra* reference genomes. Schematic representation of the methodological framework used to generate chromosome-level reference assemblies for *Porphyra dioica* and *Porphyra linearis*. The workflow encompasses: **(1)** biological material preparation (black/top panel), involving the induction of zygotospore release and establishment of homogeneous unialgal conchocelis cultures; **(2)** genome survey (blue track), assessment of ploidy level and genome size estimation via flow cytometry and k-mer profiling; **(3)** data acquisition and assembly, parallel integration of ONT long-reads (green track) and Hi-C proximity ligation (yellow track) for *de novo* assembly contig generation, scaffolding, and iterative manual curation; **(4)** repeat landscape characterization followed by structural and functional annotation (pink track) incorporating information on the transcriptome (RNA-seq) and a curated orthologous set of red algal proteins (purple tracks); and **(5)** quality assessment of the final assembly (grey panel). The photograph of the female blade is a scan of a *P. dioica* herbarium specimen (BR5010096988989V) hosted at the Meise Botanic Garden (botanicalcollections.be).

The availability of these reference genomes provides a basis to unravel the genomic determinants of physiological resilience, elucidate evolutionary patterns within the Atlantic basin, and facilitate comparative analysis of Atlantic-Pacific divergence (the European *Porphyra* vs. the Asian *Pyropia*). These resources will also accelerate future selective breeding strategies of native strains for a sustainable European aquaculture industry.

## Methods

### Establishment of unialgal conchocelis cultures

The diploid sporophytic phase (known as conchocelis, 2n) of *Porphyra dioica* and *P. linearis* was initiated from zygotospores released by wild, fertile haploid gametophytes (blade phase, n) (Fig. 1). Fertilized female gametophytes were sampled from the high-to mid-intertidal zones in Ostend, Belgium (51.239972° N, 2.925750° E; 29 January 2024) and Goes, the Netherlands (51.540776° N, 3.929514° E; 18 March 2024). Taxonomic identity for *P. dioica* (strain OSTW111) was validated on the collected gametophyte tissue via RFLP analysis of the *rbc*L gene, amplified with primers F67 and rbc-spc and digested with *Hae*III, with fragment profiles resolved by capillary electrophoresis as previously described by Teasdale *et al*. ^20^. For *P. linearis* (strain GOES6), identity was confirmed through full-length *rbcL* Sanger sequencing of the established conchocelis culture^21^.

To initiate the cultures, ∼1 cm^2^ sections of mature female tissue were rinsed in filtered, sterilized natural seawater and gently brushed to remove epiphytes, surface debris and other contaminants. Cleaned thalli were then wrapped in a moist paper towel and incubated overnight at 4°C in darkness. Zygotospore release was subsequently induced by re-immersion in modified Provasoli Enriched Seawater (mPES, 10 mL L^-1^) supplemented with GeO_2_ (1 mL L^-1^) to inhibit diatom proliferation^12,22^. Following zygotospore germination, conchocelis filaments were clonally propagated by fragmentation to generate unialgal cultures. Cultures were maintained at 15°C under a 12:12 h light:dark photoperiod at 30 μmol photons m^−2^ s^−1^, with biweekly medium renewal, and are currently maintained in the University of Ghent Culture Collection (UGCC0510 and UGCC0511) as the source material for downstream genomic and transcriptomic sequencing.

### Nuclei isolation, HMW genomic DNA extraction and ONT sequencing

To minimize microbial contamination and deplete high-copy organellar DNA, conchocelis thalli were subjected to a multi-stage decontamination and nuclei enrichment protocol. A sufficient amount of biomass was incubated in the dark for 48 hours with activated charcoal to reduce surface-associated bacteria. Subsequently, thalli were mechanically cleaned via surface abrasion by mixing the biomass with quartz sand and vortexing at 4,500 rpm using a Precellys Evolution Touch homogenizer (Bertin Technologies, Montigny-le-Bretonneux, France)^7^. The resulting homogenates were washed six times with sterile seawater to eliminate residual polysaccharides and microbial cells, flash-frozen in liquid nitrogen, and stored at -80 °C.

Nuclear enrichment was performed using a CelLytic™ PN Isolation/Extraction protocol (Sigma-Aldrich, St. Louis, MO, USA; Product No. CELLYTPN1), with modifications optimized for macroalgae. A semi-purified enrichment approach was specifically selected to maximize nuclear recovery, as high-purity pilot isolations yielded nearly no DNA for downstream sequencing. Frozen tissue (∼ 2-3 g) was pulverized in liquid nitrogen and resuspended in 10 mL of 1x Nuclei Isolation Buffer (NIB), supplemented with 0.7 M mannitol for algal osmotic stability and 1 mM DTT to prevent oxidative degradation. The suspension was filtered through two Miracloth layers (Merck, Hoeilaart, Belgium), sonicated and centrifuged at 2000 g for 10 minutes. The resulting pellet was resuspended in NIBA buffer (NIB supplemented with cOmplete™ Protease Inhibitor Cocktail; Merck, Hoeilaart, Belgium). To facilitate nuclear release while solubilizing cytoplasmic contaminants, 0.1% (v/v) Triton X-100 was added, followed by gentle agitation at 4 °C for 30 minutes. Semi-purified nuclei were obtained by layering the suspension over a 1.8 M sucrose solution and centrifuging at 12000 g for 10 minutes. The recovery was finalized with repeated NIBA washes to ensure the removal of residual cytoplasmic and organellar material.

The final purified nuclear pellets were immediately processed for HMW genomic DNA extraction following a modified CTAB-based protocol developed by Feng *et al*. ^23^. Briefly, nuclei were homogenized in CTAB lysis buffer (2% CTAB, 100 mM Tris–HCl pH 8.0, 25 mM EDTA, 2.5 M NaCl, 1% polyvinylpyrrolidone, 1% β-mercaptoethanol), incubated at 50 °C with gentle mixing, and subjected to sequential chloroform:isoamyl alcohol extractions and RNase treatment. DNA was precipitated with isopropanol, washed with 70% ethanol and resuspended in 50 μL of TE buffer. The resulting HMW DNA retained high integrity and quality suitable for long-read sequencing, as validated by NanoDrop Ultra (Fisher Scientific, Waltham, MA, USA) and 1% agarose gel electrophoresis. To further enrich long fragments and remove short-read contamination, the extracted DNA underwent a Short Read Eliminator (SRE) cleanup (PacBio, Menlo Park, CA, USA) in accordance with the manufacturer’s instructions. Final validation was performed using a 5200 Fragment Analyzer (Agilent, Zaventem, Belgium) prior to library preparation.

PCR-free ligation libraries were prepared using the Ligation Sequencing Kit V14 (SQK-LSK114; Oxford Nanopore Technologies, Oxford, UK) with concentrations monitored via Qubit 2.0 (Fisher Scientific, Waltham, MA, USA). Sequencing was performed on MinION Mk1B devices (Oxford Nanopore Technologies) using R10.4.1 (FLO-MIN114) flow cells. Each run was configured with a 72-hour duration and a pore scan frequency of 1.5 hours. Basecalling was executed using Dorado (v7.6.7) via MinKNOW (v24.11.8) in super-accurate mode with a minimum quality filter of Q ≥10. For *P. dioica*, the sequencing runs yielded 8.68 Gb of passed data (1.09 million reads; approximately 58× coverage), with a read N50 of 12.54 kb. For *P. linearis*, the generated passed data totalled 17.67 Gb (2.37 million reads; approximately 177× coverage) with a read N50 of 12.22 kb.

### Hi-C library preparation and sequencing

To achieve chromosome-scale scaffolding of the genome assemblies and resolve the three-dimensional chromatin interaction landscape, Hi-C libraries were constructed for both species. This technology allows for the orientation and anchoring of contigs by measuring the frequency of genome-wide pairwise physical contacts^24^. For each species, conchocelis tissue was fixed in a 2% formaldehyde solution in sterilized natural seawater for 60 minutes to crosslink protein-DNA and DNA-DNA interactions. The fixation was quenched by the addition of glycine for 5 minutes. Semi-purified nuclei were isolated using the Cellytic PN Isolation/Extraction Kit (Sigma-Aldrich, St. Louis, MO, USA). Hi-C libraries were generated using the Arima High Coverage HiC Kit (Arima Genomics, Carlsbad, CA, USA; A410110) following the manufacturer’s specifications. Crosslinked chromatin was digested, and the resulting 5′ overhangs were repaired with biotinylated residues, followed by *in situ* proximity ligation. The isolated proximally-ligated DNA was mechanically fragmented using a Covaris E220 Evolution ultrasonicator (Covaris Ltd, Brighton, UK) (175 W, 10% duty cycle, 200 cycles/burst) to achieve an average fragment size of 400 bp (range 200–600 bp). Fragmented DNA was size-selected using AMPure XP beads and enriched for biotinylated ligation junctions using streptavidin-coated magnetic beads. Following end-repair and adapter ligation, the libraries were amplified via 12 PCR cycles and subjected to a final purification step. Final library yields and fragment size distributions were verified using a High Sensitivity DNA kit on an Agilent 2100 Bioanalyzer (Agilent Technologies, CA, USA), resulting in 575.05 pg/µL for *P. dioica* and 621.30 pg/µL for *P. linearis*. Sequencing was performed on a NextSeq 2000 Illumina platform using a 150 bp paired-end strategy (PE150). This yielded a total of 6.29 Gb of raw data (41.7 million reads; approximately 41.9× coverage) for *P. dioica* and 11.40 Gb of raw data (75.5 million reads; approximately 114.0× coverage) for *P. linearis*. Raw data were processed with fastp (v0.23.4)^25^ to remove adapter sequences and low-quality reads. Filtering was performed using the --*detect_adapter_for_pe* flag, which effectively identified and trimmed Illumina TruSeq adapters. Ultimately, 5.98 Gb (Q30 92.4%) and 10.74 Gb (Q30 92.0%) of clean Hi-C data were retained for *P. dioica* and *P. linearis*, respectively. These high-quality datasets were subsequently utilized for chromosome-scale scaffolding and the generation of chromatin contact maps.

### RNA extraction and transcriptome sequencing (RNA-seq)

To facilitate genome annotation, total RNA was extracted from conchocelis cultures exposed to various abiotic conditions. By additionally inducing stress-responsive and metabolic genes, we aimed to capture the broadest possible representation of the transcriptome. The chosen treatments encompassed four primary physiological challenges: 1) thermal extremes, consisting of incubation at supra-optimal (21 °C) and sub-optimal (4 °C) regimes to trigger heat- and cold-shock responses; 2) hyposaline conditions, where cultures were transitioned to low-salinity media (17 ppt); 3) light attenuation, involving exposure to diminished irradiance (20 μmol photons m^−2^ s^−1^) or complete darkness; and 4) emersion-induced desiccation, achieved through a 1.5-hour period of physical dehydration to simulate the intense physiological demands of intertidal exposure. Unless otherwise noted, these stressors were applied over a 48-hour duration to stimulate a broad transcriptional response, with a parallel control group maintained under standard laboratory conditions (15°C; 12:12 h photoperiod; 30 μmol photons m^−2^ s^−1^). Following these treatments, biomass from all conditions was pooled to ensure comprehensive representation of the inducible transcriptome. Total RNA was then extracted using the RNeasy® Plant Mini Kit (Qiagen, Hilden, Germany), complying with the brand’s specifications. The resulting total RNA was confirmed to be of sufficient integrity and quality for downstream applications, following assessment on a NanoDrop Ultra spectrophotometer and a 5200 Fragment Analyzer using the RNA Kit (15 nt) (Agilent, Zaventem, Belgium).

Library preparation and high-throughput sequencing were performed by Novogene Co., Ltd. (Cambridge, UK). To specifically capture the coding fraction of the transcriptome, messenger RNA (mRNA) was purified from total RNA using poly-T oligo-attached magnetic beads. Following fragmentation, cDNA was synthesized using random hexamer primers, followed by end-repair, A-tailing, and adapter ligation. Quantified libraries were sequenced on the Illumina NovaSeq 6000 platform (PE150), generating approximately 5.7 Gb (38.0 million reads) and 5.5 Gb (36.5 million reads) of raw data for *P. dioica* and *P. linearis*, respectively. To ensure maximum accuracy for gene modelling, raw reads were processed with fastp (v0.23.4)^25^ to remove adapters and low-quality sequences. This processing substantially improved the data quality for both species: for *P. dioica*, the Q30 score increased from 93.81% to 95.04%, and for *P. linearis*, the Q30 score increased from 93.57% to 94.90%. Post-filtering quality control confirmed high reliability for both species, with an average base error rate of 0.01% and an effective read rate exceeding 97.70%, providing a high-confidence dataset for downstream *ab initio* gene prediction and transcript-supported modelling.

### Genome survey

#### Genome Size Estimation via Flow Cytometry

Prior to assembly, the nuclear DNA content of both species was quantified using flow cytometry (Fig. 2). For all analyses, conchocelis tissue, representing the filamentous diploid sporophyte phase, was thoroughly rinsed with Milli-Q water and gently blotted dry before processing. Nuclear suspensions for *P. dioica* were prepared at the FLOWer Lab (University of Coimbra, Portugal) by co-chopping approximately 20 mg of algal tissue with the internal reference standard, *Raphanus sativus* (2C DNA content = 0.98 pg; 2C genome size = 956 Mbp)^26^ in 1 mL of WPB buffer as described in Varela-Álvarez *et al*. ^13^. Tissue was then filtered through a 50 μm nylon mesh and stained with 50 μg/mL of propidium iodide (PI) and 50 μg/mL of RNase to prevent double-stranded RNA interference. Samples were analyzed within 5 minutes of preparation using a Partec CyFlow Space flow cytometer (Partec GmbH, Görlitz, Germany) equipped with a green solid-state laser for PI excitation. For *P. linearis*, nuclear suspensions were prepared at the Flemish Institute for Biotechnology (VIB) - UGent Center for Plant Systems Biology (Ghent, Belgium) by co-chopping algal tissue with *Solanum lycopersicum* (2C DNA content = 1.72 pg; 2C genome size = 1682 Mbp)^26^ in 500 µL of ice-cold LB01 buffer^27^. An additional 500 µL of LB01 was added, and the homogenate was stored on ice. The homogenate was then filtered through a 42 μm nylon mesh and incubated at 4°C for 1h. Subsequently, the upper 200 μL of the supernatant was extracted and treated with 550 μl of a staining solution containing LB01 lysis buffer with 50 μg mL^−1^ PI, 50 μg mL^−1^ RNase IIA, and 2 μl mL^−1^ β-mercaptoethanol. Stained samples were incubated in the dark at 4°C for 10 minutes and analysed using an Attune NxT flow cytometer (Fisher Scientific, Waltham, MA, USA) equipped with a 561 nm yellow excitation laser (50 mW) and a 620/15 emission filter (YL2 channel).

**Figure 2.**
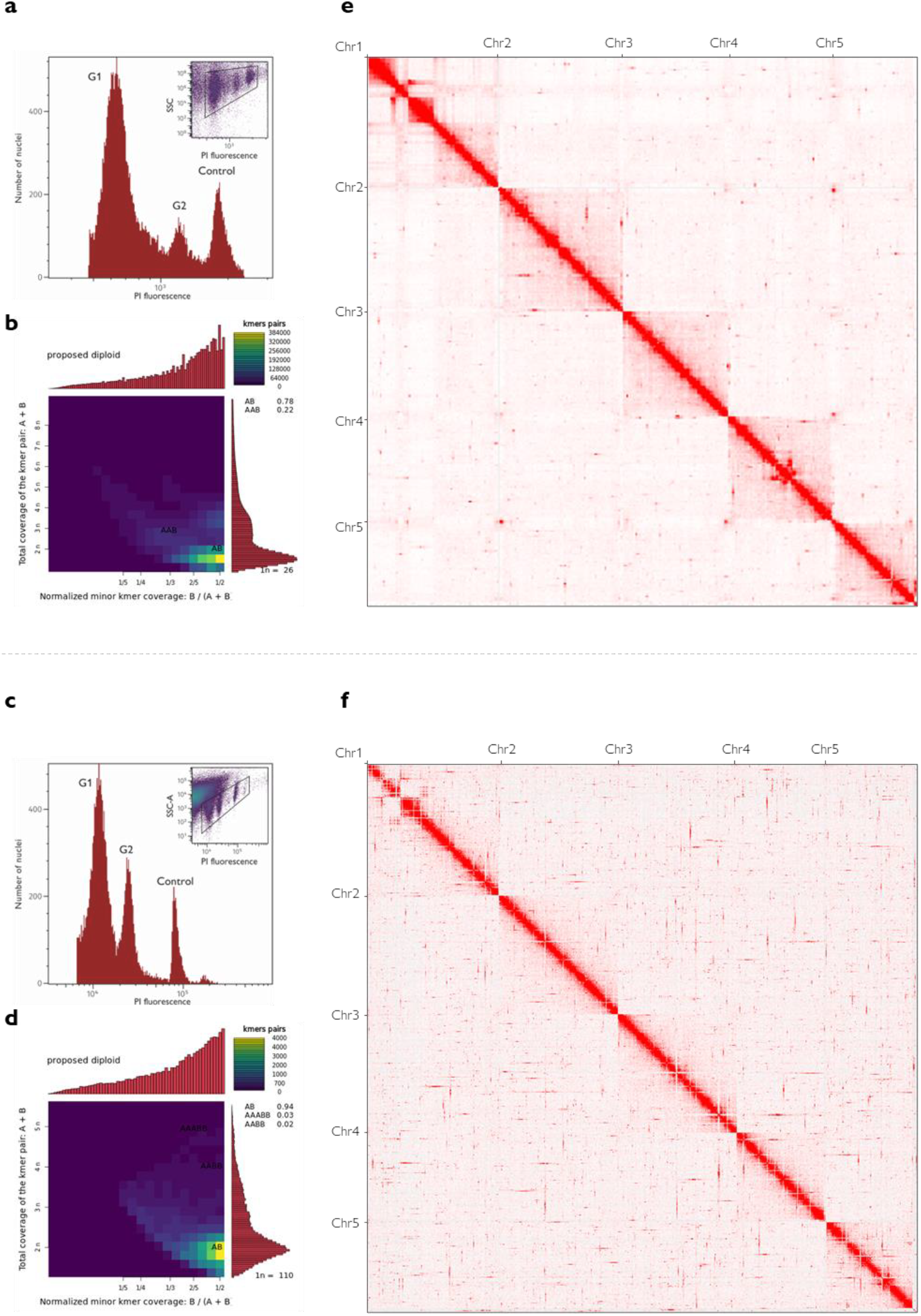
Genome survey and chromosome-scale validation of *Porphyra dioica* and *Porphyra linearis*. Panels in the top and bottom rows correspond *to P. dioica* and *P. linearis*, respectively. **(a, c)** Flow cytometry histograms of propidium iodide-stained conchocelis nuclei, showing G1 and G2 peaks alongside the internal reference standard as a control; insets display the gating strategy (SSC or SSC-A vs PI intensity) used to isolate target nuclei from background debris. **(b, d)** Smudgeplot inference of ploidy level based on heterozygous k-mer pairs from raw ONT reads. The dominant “AB” signal confirms a diploid (2n) state for both species, determined by the ratio of k-mer coverage sums (CovA + CovB) to relative coverage (CovB / (CovA + CovB)). **(e, f)** Normalized genome-wide chromatin interaction heatmaps for the curated assemblies of *P. dioica* (∼150 Mbp) and *P. linearis* (∼95 Mbp). The axes denote five pseudo-chromosomes (n=5) per species, with colour intensity representing contact frequency. The sharp diagonal signals and distinct off-diagonal blocks confirm high-contiguity scaffolding and structural integrity.

For both species data were analysed with the FlowJo v11 software (BD Biosciences). Nuclei were gated by plotting side scatter (SSC) against PI intensity for *P. dioica* and side scatter area (SSC-A) against PI intensity for *P. linearis* to effectively isolate the target G1 populations from background debris (Figs. 2a, 2c). Only gates containing more than 1000 nuclei in the G1 peaks were used for reliable genome size calculations (Sliwinska *et al*. 2022). Genome sizes (in Mbp) of each sample were determined by applying the following formula to the median PI value of G1: Mbp_sample = (Mbp_control / Median_control) · Median_sample. Based on this methodology, the estimated diploid genome sizes were 332.64 Mbp for *P. dioica* and 235.40 Mbp for *P. linearis*.

#### K-mer based genome profiling

To supplement the flow cytometry data, a computational assessment of genome size, heterozygosity, repeat content, and ploidy level was conducted using k-mer profiling of the raw ONT reads. K-mer frequency distributions were generated using Jellyfish (v2.3.1)^28^ (k = 21) and analysed with GenomeScope (v1.0 and v2.0)^29,30^. For *P. dioica*, profiling with GenomesScope (v2.0) indicated a most likely haploid genome size of 146.85 Mbp (model fit: 95.60%), with a heterozygosity rate of 1.84% and a repetitive fraction of approximately 77.14 Mbp (52.53% of the estimated haploid span). For *P. linearis*, GenomeScope (v1.0) yielded a most likely haploid genome size of 98.09 Mbp (model fit: 92.23%), characterized by a notably high heterozygosity rate of 3.61% and a substantial repetitive fraction accounting for 63.09 Mbp (∼64.32% of the estimated haploid span). To ensure the accuracy of the ploidy estimation, k-mer counts were filtered using the Smudgeplot (v0.5.3)^30^ *cutoff* tool to exclude low-frequency sequencing noise (*L*) and excessively high-frequency organellar or repetitive DNA (*U*) (*L*=15, *U*=1800 for *P. dioica*; *L*=76, *U*=320 for *P. linearis*). Final validation and visualization were performed using the Smudgeplot (v0.2.5+galaxy3) on the European Galaxy web portal (https://usegalaxy.eu/)^31^, which confirmed the diploid (AB) state as the most probable ploidy level for both species (Figs. 2b, 2d).

### *De novo* genome assembly and chromosome construction

Chromosome-level *de novo* reference genomes were generated for *P. dioica* and *P. linearis* by anchoring ONT long-reads to Hi-C chromatin interaction maps.

#### Initial contig-level assembly and polishing

To identify the optimal assembly strategy for the *Porphyra* datasets, several long-read assemblers were benchmarked. Performance was evaluated based on the recovery of conserved orthologs (BUSCO v6.0.0)^32^ and overall genomic statistics (QUAST v5.2.0)^33^. Based on these metrics, the Flye (v2.9.6) assembler^34^ (utilizing four rounds of internal iterative polishing; --*iterations* 4), followed by three sequential rounds of polishing with Racon (v1.5.0)^35^, was selected as the best-performing approach. To maximize sequence accuracy, base-level consensus refinement was achieved using Medaka (v1.12.1)^36^ with the high-accuracy model r1041_e82_400bps_sup_v5.0.0, specifically optimized for the ONT R10.4.1 chemistry.

At this stage, the primary assembly for *P. dioica* reached a total span of 164.86 Mbp across 2,427 contigs, featuring a largest contig of 4.31 Mbp and a GC content of 65.59% (N50 = 135.49 kb). In comparison, the *P. linearis* assembly reached a total span of 183.28 Mbp across 1,451 contigs, with a largest contig of 6.63 Mbp and a GC content of 59.87% (N50 = 315.41 kb). Both assemblies were characterized by a total absence of gaps, providing a continuous and robust basis for subsequent chromosome-scale scaffolding.

#### Chromosome-scale scaffolding and manual curation

Chromosome-scale architecture was achieved by anchoring the consensus-refined contigs using Hi-C proximity ligation data. Cleaned Hi-C reads were mapped to the primary assemblies using bwa (v0.7.17)^37^ with the -*5SP* option. The resulting alignments were filtered with samblaster (v0.1.26)^38^ and SAMtools (v1.17)^39^ to retain high-confidence chromatin interactions (bitmask filter of -*F 3340*, effectively excluding unmapped reads, non-primary alignments, and duplicate sequences).

Automated scaffolding was performed using the HapHiC (v1.0.7) pipeline^24^. To ensure the assemblies conformed to the known karyotypic organization of the Bangiales, we targeted a structure of five chromosomes (n=5). This count is consistent with previous cytogenetic reports for both target species: n=5 for *P. dioica*^40^ and n=5 for *P. linearis*^41^. This karyotype was empirically validated using the HapHiC -- *quick_view* utility, which generated a preliminary ordering of contigs without a priori input regarding chromosome number. The resulting visualizations revealed five distinct interaction territories for both species, confirming that the physical interaction data independently resolved the expected chromosome count. Final scaffolding was then executed incorporating the motifs corresponding to the Arima-HiC multi-enzyme cocktail utilized in the library preparation (--*GATC, GANTC, CTNAG, TTAA*) along with two rounds of iterative orientation and ordering (--*correct_nrounds 2*). The resulting scaffolds were manually curated in Juicebox (v1.8.8)^42^. Normalized (*balanced* mode) Hi-C contact maps were used to visualize interaction frequencies (Figs. 2e, 2f), allowing for the correction of misjoins, resolution of translocations, and reorientation of scaffolds. We utilized the HapHiC *juicer post* utility to reconcile these manual edits and generate the finalized chromosome-level FASTA sequences.

#### Purification of the nuclear genome

Beyond structural scaffolding, the integration of Hi-C data provided a robust mechanism for discriminating the target nuclear genomes from organellar signals and associated microbial communities inherent in algal cultures^43^. Within this context, three independent lines of evidence were employed to isolate non-nuclear scaffolds and ensure the generation of a high-purity reference assembly:

1. *Hi-C interaction topology*: Scaffolds lacking significant interaction signals with the five primary nuclear territories — appearing as isolated clusters or “noise” off the diagonal of the contact map — were identified as potential exogenous contaminants and excluded.
2. *Automated taxonomic classification*: Tiara (v1.0.2)^44^ was utilized to classify all scaffolds into “eukarya”, “bacteria”, “organelle”, or “unknown” categories. Only sequences identified as eukaryotic or consistent with the *Porphyra* nuclear signal were retained.
3. *GC content and coverage profiles*: Using bioawk^45^ and SAMtools (v1.21)^39^, we calculated the GC content and sequencing depth per scaffold. Nuclear sequences were retained based on their characteristic enrichment within the 55–75% GC range and a stable coverage profile (100– 200x). Conversely, scaffolds with low GC content and anomalous coverage — representing either high-depth organellar DNA or low-depth bacterial contaminants — were purged.

This filtering approach successfully removed approximately 14 Mbp (∼8.5%) and 87 Mbp (∼47.5%) of non-target sequences from the *P. dioica* and *P. linearis* assemblies, respectively. The finalized, purified assemblies reached 150.44 Mbp for *P. dioica* and 95.22 Mbp for *P. linearis*. Both genomes were successfully resolved into five nuclear chromosomes (n=5), accounting for nearly 100% of the total assembly span. A small fraction of unplaced nuclear contigs exhibiting inter-chromosomal interactions were sequestered as unanchored fragments; consequently, functional annotation was restricted to the five high-confidence chromosomal scaffolds. In *P. dioica*, the largest chromosome reached 36.54 Mbp with an assembly N50 of 29.04 Mbp and an overall GC content of 66.04%. Comparatively, in *P. linearis*, the largest chromosome reached 22.99 Mbp with an N50 of 20.87 Mbp and an overall GC content of 67.73 (Fig. 3). Chromosomes were numbered by descending sequence length (Chr1–Chr5), with Hi-C heatmaps revealing distinct structural heterogeneity along the diagonal of Chr1 in both species (Figs. 2e, 2f).

**Figure 3.**
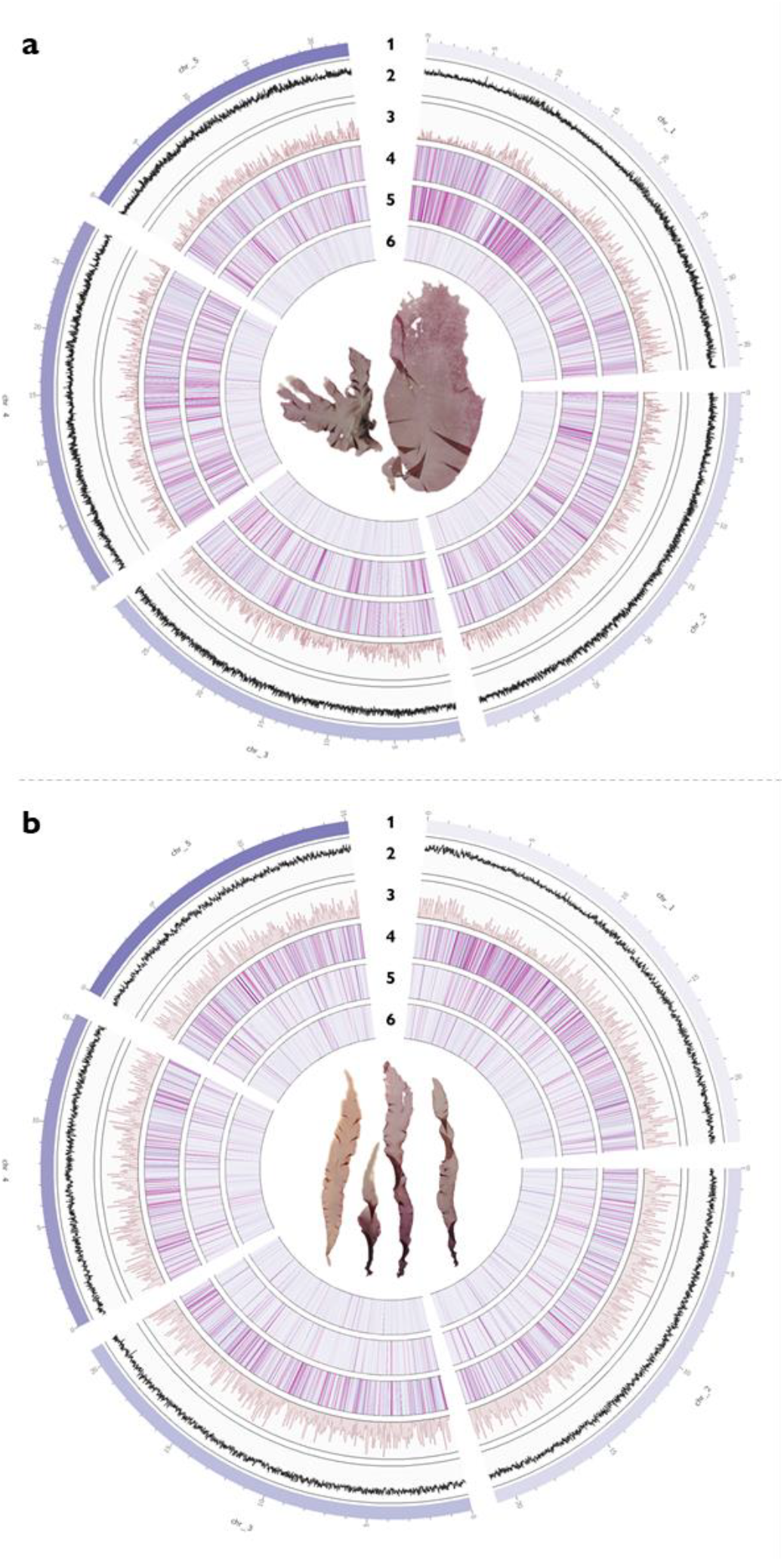
Comparative chromosome-level genomic architecture of *Porphyra dioica* and *Porphyra linearis*. Circos plots illustrate the spatial distribution of genomic features across the five nuclear chromosomes of **(a)** *P. dioica* and **(b)** *P. linearis*. From outside to inside, tracks represent: **(1)** Chromosome ideograms scaled in megabases (Mb); **(2)** guanine-cytosine (GC) content in 10 kb windows; **(3)** protein-coding gene density in 50 kb windows, representing primary isoforms retained after AGAT filtering; **(4)** unclassified repetitive elements, comprising 22.36% and 21.80% of the *P. dioica* and *P. linearis* genomes, respectively; **(5)** Long Terminal Repeat (LTR) retrotransposon density; and **(6)** DNA transposon content. Tracks 4–6 constitute the three major repetitive classes characterized in this study, plotted in 10 kb windows. Central photographs are scans of representative herbarium specimens hosted at Meise Botanic Garden: BR5010096993846V and BR5010096988989V (*P. dioica*); BR5010096723719V and BR5010105785561V (*P. linearis*). Detailed repeat landscapes and annotation statistics are provided in Table 1 and Table 2.

### Structural, functional and repeat annotation

#### Repeat landscape characterization and masking

Prior to the prediction of protein-coding genes, repetitive elements and transposable elements (TEs) were identified and soft-masked across the chromosome-level assemblies of each species. A two-step annotation framework, adapted from Petroll *et al*. ^46^, was employed.

**Table 1.**
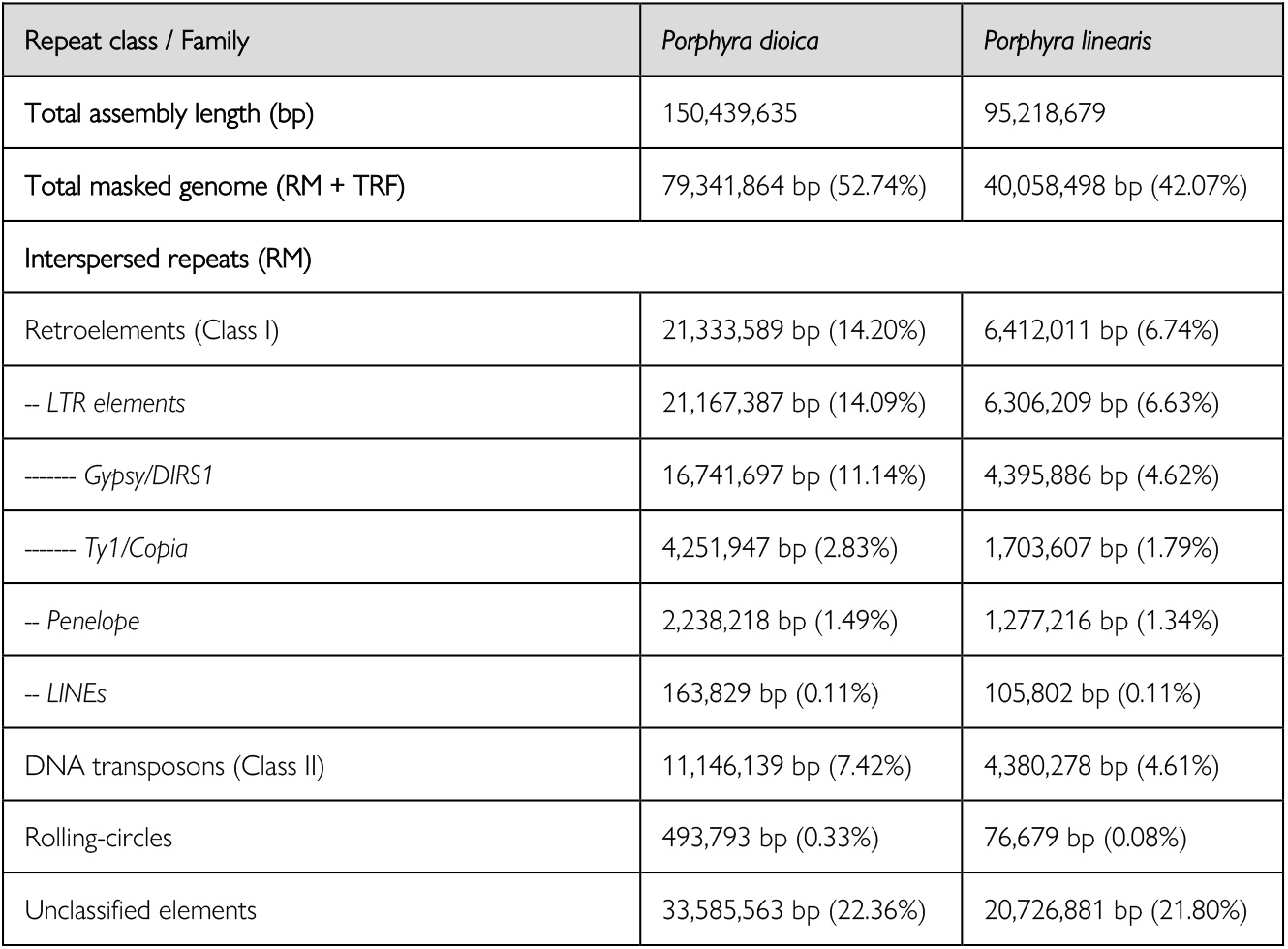
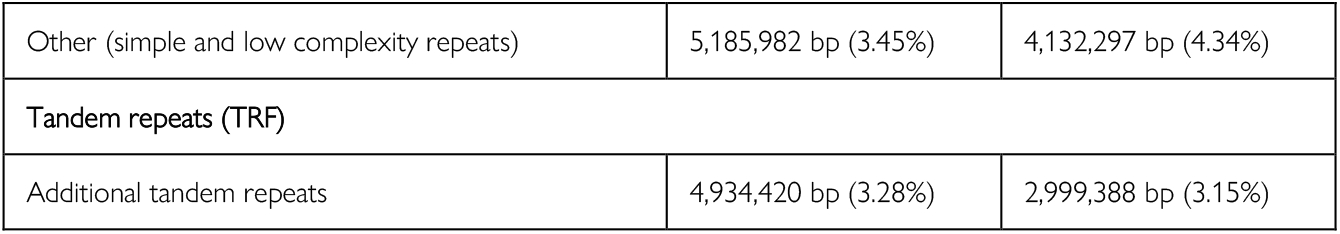
Summary of repetitive elements identified in the chromosome-level assemblies of *Porphyra dioica* and *Porphyra linearis*. Interspersed elements were identified via RepeatModeler (RM), while additional tandem arrays were supplemented via Tandem Repeats Finder (TRF).

| Repeat class / Family | *Porphyra dioica* | *Porphyra linearis* |
| --- | --- | --- |
| Total assembly length (bp) | 150,439,635 | 95,218,679 |
| Total masked genome (RM + TRF) | 79,341,864 bp (52.74%) | 40,058,498 bp (42.07%) |
| Interspersed repeats (RM) |  |  |
| Retroelements (Class I) | 21,333,589 bp (14.20%) | 6,412,011 bp (6.74%) |
| – LTR elements | 21,167,387 bp (14.09%) | 6,306,209 bp (6.63%) |
| ----- Gypsy/DIRS1 | 16,741,697 bp (11.14%) | 4,395,886 bp (4.62%) |
| ----- Ty1/Copia | 4,251,947 bp (2.83%) | 1,703,607 bp (1.79%) |
| – Penelope | 2,238,218 bp (1.49%) | 1,277,216 bp (1.34%) |
| – LINEs | 163,829 bp (0.11%) | 105,802 bp (0.11%) |
| DNA transposons (Class II) | 11,146,139 bp (7.42%) | 4,380,278 bp (4.61%) |
| Rolling-circles | 493,793 bp (0.33%) | 76,679 bp (0.08%) |
| Unclassified elements | 33,585,563 bp (22.36%) | 20,726,881 bp (21.80%) |
| Other (simple and low complexity repeats) | 5,185,982 bp (3.45%) | 4,132,297 bp (4.34%) |
| **Tandem repeats (TRF)** |  |  |
| Additional tandem repeats | 4,934,420 bp (3.28%) | 2,999,388 bp (3.15%) |

**Table 2.** Summary of structural and functional annotation statistics for the Porphyra dioica and Porphyra linearis genomes. Protein-coding statistics represent the final non-redundant gene sets following AGAT refinement. Values represent total counts across the five nuclear chromosomes.

| Feature | *Porphyra dioica* | *Porphyra linearis* |
| --- | --- | --- |
| **Structural annotation** |  |  |
| Total predicted protein-coding genes | 12,548 | 10,382 |
| Total mRNA transcripts | 13,845 | 10,609 |
| Mean gene/CDS length (bp) | 1,346 / 1,260 | 1,386 / 1,286 |
| Mean exons per transcript | 1.3 | 1.3 |
| Single-exon genes (%) | 80.3% | 78.5% |
| **Functional assignment of protein-coding genes** |  |  |
| Genes with eggNOG/COG (Total/Known*) | 4,564 / 3,455 | 3,811 / 2,889 |
| Genes with InterProScan domains | 6,933 | 5,908 |
| Genes with assigned GO terms | 3,097 | 2,700 |
| Genes with KEGG (KO) terms | 4,252 | 3,643 |
| Total Functional Annotation Rate (%) | 57.4% | 59.8% |
| **Non-coding RNA (ncRNA)** |  |  |
| Total tRNAs (Standard/Pseudo) | 184 / 36 | 185 / 23 |
| Total rRNA loci (5S / 45S components) | 75 / 2-4 | 68 / 3-5 |

\*Known indicates A–Q categories, excluding category S.

To characterize interspersed repeats, species-specific TE libraries were constructed using RepeatModeler (v2.0.5)^47^, which performed *de novo* repeat discovery and taxonomic classification against the Dfam database. The resulting custom libraries were subsequently deployed as input for RepeatMasker (v4.1.5)^48^ to soft-mask the genomes (cutoff score = 225; parameters: -xsmall, -gff, -excln). In line with the genomic features observed in related Bangiales^6-8,19^, we identified a substantial amount of repetitive sequences comprising 49.46% and 38.92% of the *P. dioica* and *P. linearis* genomes, respectively. Within this repetitive fraction, both landscapes were dominated by unclassified elements, followed by long terminal repeat (LTR) retrotransposons and DNA transposons (Table 1; Fig. 3).

To resolve tandem arrays, satellite and microsatellite elements were further identified across the unmasked assemblies utilizing Tandem Repeats Finder (TRF, v4.09.1)^49^ with optimized scoring parameters (match = 2, mismatch = 7, delta = 7, PM = 80, PI = 10, minscore = 50, maxperiod = 2000). The resulting TRF coordinates were consolidated using BEDTools (v2.31.1)^50^ and incorporated into the RepeatMasker-processed genomes via the *maskfasta* utility (with the -soft parameter)^50^. As a result, this integration successfully captured an additional 3.28% and 3.15% of tandem sequence in *P. dioica* and *P. linearis*, respectively, yielding cumulative soft-masked fractions of 52.74% and 42.07% (Table 1).

#### Structural gene prediction

Following the soft-masking of repetitive elements, we implemented the BRAKER3 (v3.0.8+galaxy2) pipeline^51^ through the European Galaxy platform (https://usegalaxy.eu/)^31^ to perform automated, evidence-driven *ab initio* gene model prediction. This hybrid framework integrated species-specific transcriptomic data with a curated orthologous protein database.

To provide transcriptomic support, trimmed RNA-seq short-reads were mapped to the respective soft-masked assemblies using HISAT2 (v2.2.1)^52^ with the *--dta* and *RF* strand-specific options to provide precise splice-junction evidence for downstream training. In parallel, a curated orthologous set of protein sequences was derived from five published Rhodophyta proteomes, namely: *Porphyra umbilicalis*^53^, *Neopyropia yezoensis*^54^, *Pyropia vietnamensis*^55^, *Neoporphyra haitanensis* (as *Py. haitanensis* PH40)^*19*^, and *Chondrus crispus*^56^. For this, orthologous genes and orthogroups were identified using OrthoFinder (v2.5.5+galaxy1)^57^, yielding an initial set of 9,323 orthogroups. In order to prioritize biologically robust signals, a stringent occupancy filter was applied, retaining only those orthogroups present in at least 80% of the reference species. This strategy resulted in a refined set of 4,113 conserved orthogroups, comprising 2,063 groups shared across all five species and 2,050 present in four, which were merged into a single amino acid database (13Mb) to serve as homology evidence for gene prediction.

By integrating RNA-seq alignments and the curated protein database, the BRAKER3 pipeline automatically executed in ETP mode, enabling the simultaneous training of GeneMark-ETP and AUGUSTUS with five rounds of optimization. Initial predictions identified 12,514 and 10,351 protein-coding loci for *P. dioica* and *P. linearis*, respectively. Structural analysis revealed a predominantly single-exon architecture, with multi-exon transcripts accounting for only ∼20% of the total gene set. Mapping of these loci revealed that chromosome 2 is the most gene-dense in both species, harboring 3,083 loci in *P. dioica* and 2,615 in *P. linearis* (Fig. 3, track 3). To refine these models, raw predictions were processed using Another GFF Analysis Toolkit (AGAT) (v1.4.0+galaxy1)^58^. We utilized the *agat_sp_keep_longest_isoform*.*pl* script to retain only the primary transcript per locus and resolve overlapping features, resulting in a refined final non-redundant set of 12,548 and 10,382 genes (Table 2).

#### Functional annotation

Functional characterization of the predicted protein-coding genes was performed using an integrated phylogenomic and domain-based approach facilitated by the European Galaxy server^31^. Orthology assignments and functional transfers were executed using eggNOG-mapper (v2.1.13)^59^ based on the eggNOG database (v5.0.2). Sequence searches were conducted using the DIAMOND engine in sensitive mode (--*sensmode* sensitive). This analysis provided annotations across three hierarchical levels: Gene Ontology (GO) terms for discrete biological processes, KEGG pathways for metabolic and signaling reconstruction, and Clusters of Orthologous Groups (COG) for broad evolutionary classification.

In parallel, protein domains and functional sites were identified using InterProScan (v5.59-91.0)^60^, integrating signature matches from Pfam (v35.0)^61^, PANTHER (v17.0)^62^, and SUPERFAMILY (v1.75)^63^. To ensure exhaustive local calculation and high-sensitivity detection of lineage-specific domains, the pre-calculated match lookup service was disabled. The resulting annotations were reconciled to support metabolic reconstruction and ensure the dataset is fully compatible with global genomic databases.

The annotation pipeline successfully assigned biological descriptions or conserved domains to a substantial proportion of both protein-coding genes, achieving a total functional annotation rate of 57.4% (7,202 loci) for *P. dioica* and 59.8% (6,208 loci) for *P. linearis* (Table 2). Beyond broad orthology, domain-based analysis via InterProScan identified conserved signatures in 55.3% and 56.9% of the respective gene sets, while over 25% of the total loci were successfully mapped to Gene Ontology (GO) terms. While the relative distribution of COG functional categories remained remarkably consistent between species, *P. dioica* possessed a higher absolute gene count across all functional classes, consistent with its overall genome expansion. Excluding the “Function unknown” (S) class, the functional profiles of both species were primarily dominated by genes involved in post-translational modification, protein turnover, and chaperones (Category O; 740/641 loci for *P. dioica*/*P. linearis*), replication, recombination, and repair (Category L; 590/379), translation and ribosomal structure (Category J; 403/364), and signal transduction mechanisms (Category T; 353/262).

#### Annotation of non-coding RNA genes

To complete the genomic architecture, candidate non-coding RNA (ncRNA) genes were identified across both assemblies using specialized eukaryotic models. Transfer RNAs (tRNAs) were predicted using tRNAscan-SE (v2.0.12)^64^, yielding 220 genes in *P. dioica* and 208 in *P. linearis*, with a notably high proportion of intron-containing types (35.45% and 30.28%, respectively). Ribosomal RNA (rRNA) subunits were characterized using barrnap (v0.9)^65^. *P. dioica* exhibited a robust recovery of 5S rRNA loci (75 copies) primarily organized in tandem arrays on chromosome 1, while *P. linearis* contained 68 identified 5S loci distributed across chromosomes 1 and 4. Both species maintained a conserved set of 45S cistron components (18S, 5.8S, and 28S), with 2–5 loci identified across the nuclear chromosomes (Table 2).

### Organelle genome assembly and annotation

Following the construction of the chromosome-level nuclear assembly, candidate organellar scaffolds were partitioned from the nuclear background using a multi-faceted approach that paired Tiara-based taxonomic classification, alongside characteristic GC content and sequencing depth. In both species, organellar (and bacterial) sequences were clearly distinguished by an elevated copy number (∼18- to 180-fold higher coverage) and significantly lower GC content (∼32%) relative to the nuclear genome (∼64–66%). These specific scaffolds were extracted using SeqKit (v2.8.2)^66^ and imported into Geneious Prime (v2026.0.2; https://www.geneious.com) for manual structural finalization. First, the organellar scaffolds were identified via BLAST similarity searches against reference *Porphyra purpurea* chloroplast (NC_000925.1)^67,68^ and mitochondrial (NC_002007.1)^69,70^ genomes. Overlapping terminal regions were identified to circularize each contig, resulting in complete, circular chloroplast and mitochondrial genomes (Fig. 4). Gene models were projected onto the finalized contigs using the ‘*Map to Reference*’ tool with high-sensitivity parameters, utilizing the aforementioned *P. purpurea* references. Annotation completeness was assessed using the ‘*Find ORFs*’ tool in Geneious followed by BLASTp searches^71^ against the NCBI nonredundant database (http://blast.ncbi.nlm.nih.gov/Blast.cgi). All annotations were visually inspected and adjusted where necessary. rRNA boundaries were defined by aligning sequences to a curated dataset of red algal cpDNAs and mtDNAs, while mitochondrial cox1 intron-exon boundaries were refined through sequence alignment with published orthologs. tRNA gene predictions were further validated using the tRNAscan-SE 2.0^64^ web server (https://trna.ucsc.edu/tRNAscan-SE/).

**Figure 4.**
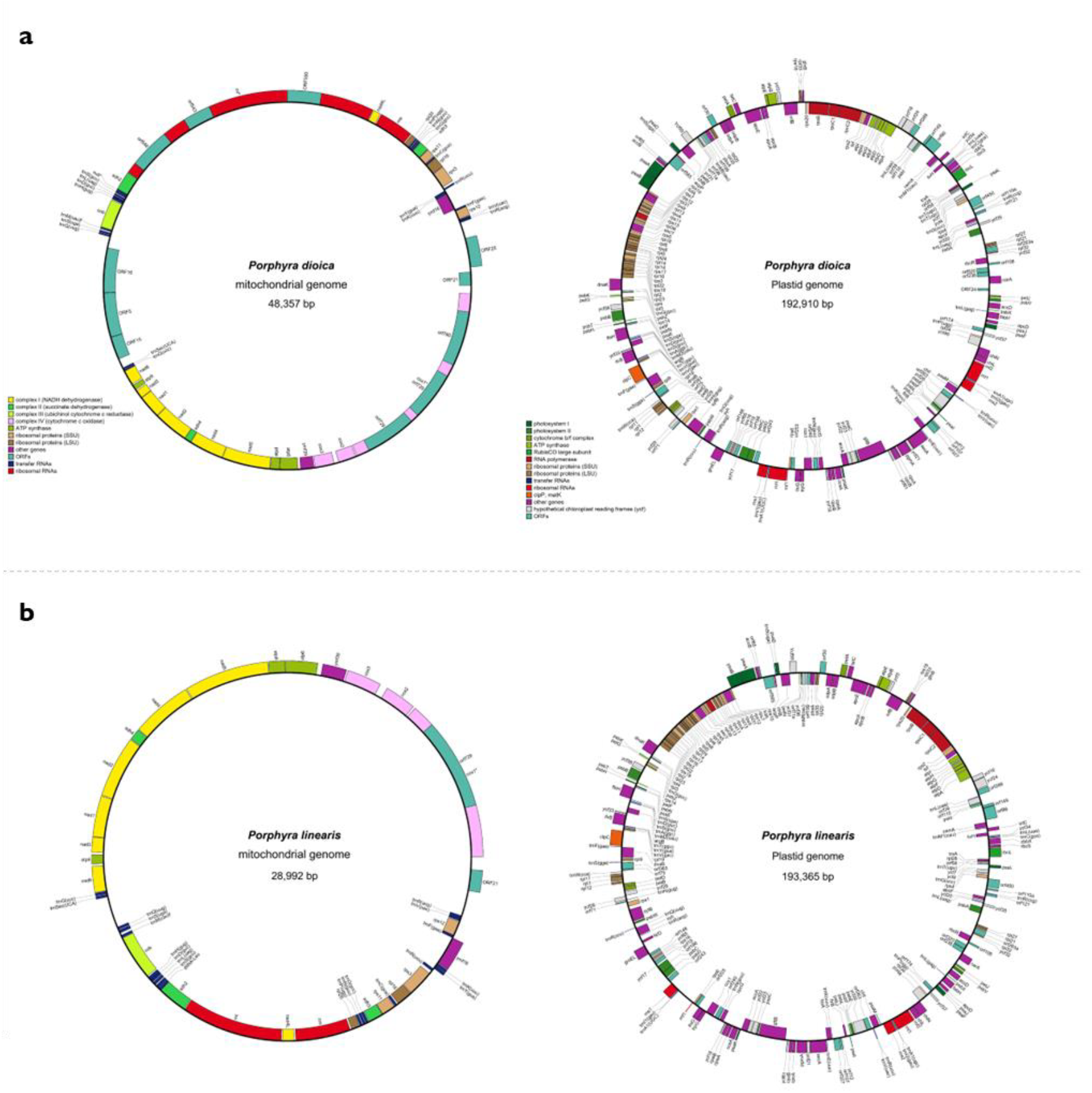
Circular maps of the mitochondrial and chloroplast genomes. Structural organization and gene content for **(a)** *Porphyra dioica* and **(b)** *Porphyra linearis*. Genes are color-coded by functional category as indicated in the legend. All maps were generated using OGDRAW^72^ (https://chlorobox.mpimp-golm.mpg.de/OGDraw.html).

## Data records

The genome assemblies and raw sequencing data generated for *Porphyra dioica* and *Porphyra linearis* are publicly available and have been deposited in the European Nucleotide Archive (ENA) under the umbrella BioProject accessions PRJEB104629 and PRJEB104630, respectively.

## Technical Validation

### Quality assessment

The structural integrity of the chromosome-level assemblies was assessed via QUAST (v5.2.0)^33^, which confirmed that both genomes are represented by five high-contiguity scaffolds (L50 = 3) corresponding to the primary nuclear chromosomes. *Porphyra dioica* (150.44 Mbp) reached an N50 of 29.04 Mbp, while *Porphyra linearis* (95.22 Mbp) yielded an N50 of 20.87 Mbp. Base composition analysis revealed overall GC contents of 66.04% and 67.73%, respectively—a compositional signature highly consistent with the characteristic GC-richness of the Bangiophyceae^7^. This attribute aligns with established benchmarks for *P. umbilicalis* (∼66%)^6^ and *Bangia fuscopurpurea* (∼64%)^73^, as well as the East Asian chromosome-level references *Neopyropia yezoensis* (∼65%)^7^ and *Neoporphyra haitanensis* (∼70%)^8^. The high-fidelity recovery of these GC-rich profiles provides orthogonal evidence for the successful isolation of target nuclear DNA and the exhaustive removal of microbial contaminants. Complementing these metrics, the physical scaffold lengths show high congruence with haploid genome sizes estimated via flow cytometry (∼166 and ∼118 Mbp, respectively) and k-mer profiling (146.85 and 98.09 Mbp), indicating a near-complete capture of the genomic span.

### Genomic and transcriptomic completeness

The completeness of the genome assemblies was benchmarked using BUSCO (v6.0.0)^32^ in genome mode. Evaluation against the rhodophyta_odb12 lineage (n=1,591) yielded completeness scores of 71.8% for *P. dioica* and 71.4% for *P. linearis*, while searches against the broader eukaryota_odb10 dataset (n=255) resulted in scores of 58.0% and 58.8%, respectively. As noted by Hanschen *et al*. ^74^, standard ortholog-based assessment tools are often not highly applicable to algal genomes due to their extraordinary phylogenetic diversity and early branching nature. Their analysis demonstrated that while BUSCO gene sets are designed to be present in over 90% of species within a group, less than 19% of the Eukaryota models actually meet this threshold across the diverse algal lineages. Consequently, our scores reflect inherent evolutionary divergence rather than assembly deficiency and are consistent with other red algal references. To further validate the representation of the expressed gene space, trimmed RNA-seq reads were re-mapped to the final assemblies using HISAT2^52^, yielding overall alignment rates of 88.24% for *P. dioica* and 92.62% for *P. linearis* and confirming that the assemblies captured the vast majority of the expected gene space.

### Chromosomal synteny and evolutionary conservation

To evaluate structural accuracy and evolutionary conservation of the generated assemblies, we performed a whole-genome synteny analysis between *P. dioica* and *P. linearis*. All-to-all protein sequence alignments were executed using BLASTp (v2.17.0)^71^ with a stringent E-value threshold of 1e^-10^ and a maximum of five target sequences per query. Collinear blocks and gene duplications were subsequently identified using the MCScanX toolkit^75^ with default parameters. The resulting syntenic relationships were visualized using the web-based tool SynVisio (https://synvisio.github.io/)^76^, revealing a robust one-to-one correspondence across all five chromosomes (Fig. 5). This high degree of chromosomal conservation reflects the close evolutionary relationship between these species^21,77^, validates the accuracy of our Hi-C-based scaffolding and provides a reliable foundation for comparative genomic studies.

**Figure 5.**
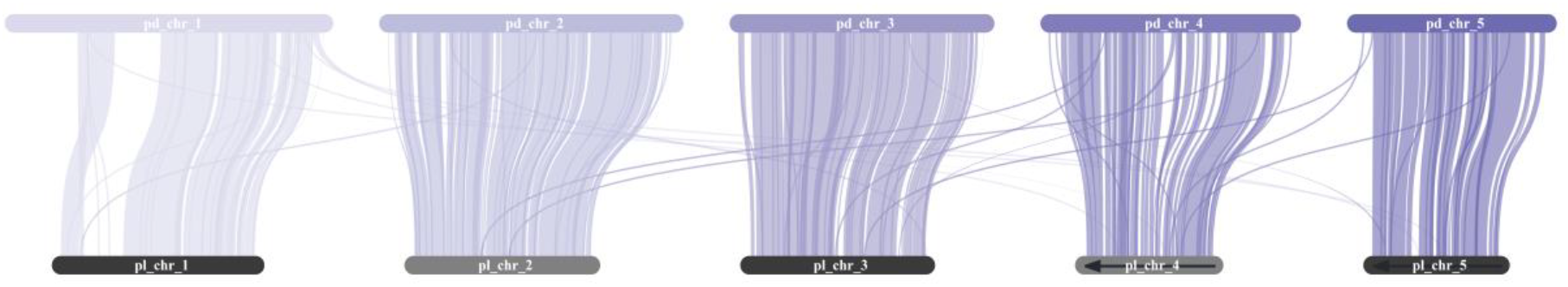
Chromosomal synteny and collinearity between *Porphyra dioica* and *Porphyra linearis*. Macrosynteny plot illustrating the structural conservation across the five nuclear chromosomes of *P. dioica* (top; purple) and *P. linearis* (bottom; grey). Connecting ribbons represent homologous gene blocks. Arrows on the *P. linearis* ideograms indicate the relative orientation of anchored scaffolds. Inter-chromosomal ribbons suggest localized rearrangements and translocations. Ideograms are scaled proportionally to their physical assembly lengths (Mbp).

## Code availability

All bioinformatics analyses in this study were conducted using established, publicly available software packages in accordance with standardized operational protocols. No custom scripts, proprietary algorithms, or non-standardized code implementations were employed for the generation or processing of the datasets. Detailed information regarding software versions and specific parameters is provided within the respective Methods sections. Unless otherwise specified, all tools were executed using default parameter settings as per the official documentation of the respective software packages. Any additional information or specific command-line implementations required to reanalyse the data reported in this study are available from the corresponding author upon request.

## Acknowledgements

This project has received funding from the European Union under the European Union’s Horizon Europe research and innovation programme under the Biodiversity, Circular Economy and Environment (REA.B.3); co-funded by the Swiss State Secretariat for Education, Research and Innovation (SERI) under contract number 24.00054 and by the UK Research and Innovation (UKRI) under the Department for Business, Energy and Industrial Strategy’s Horizon Europe Guarantee Scheme. Further funding was provided by cSBO GAME [HBC.2023.0503]. Individual financial support was provided to J.M. by the Fonds Wetenschappelijk Onderzoek (FWO) [1148826N]. We also thank Mariana Castro and João Loureiro from the FLOWer Lab, University of Coimbra, for their expertise and technical assistance in generating the flow cytometry data for *P. dioica*.

## Author contributions

**O.D.C**. and **J.K**. conceived the study, secured funding, and provided overall project supervision. **J.M**., **J.K**. and S.B. collected biological samples and established homogeneous unialgal cultures. **S.D**., **J.M** and **A.P.L**. generated the genomic and transcriptomic data, including the preparation of ONT, Hi-C, and RNA-seq libraries and sequencing. **J.M**. performed the *de novo* nuclear genome assembly, iterative polishing, Hi-C scaffolding, and manual curation. **F.L**. reconstructed and annotated the plastid and mitochondrial genomes. **J.M**. conducted the structural and functional gene annotation of the nuclear assembly and repeat landscape characterization, with support from **L.N. J.M**. performed technical validation, including k-mer profiling, ploidy validation, and assembly completeness assessments. **S.B**. and **A.V.d.V**. performed the flow cytometry for genome size estimation and provided the histograms. **J.M**. managed data curation and public repository submissions. **J.M**. drafted the manuscript and designed the figures. **S.V**. and **A.P.L**. contributed to the critical review of the manuscript. All authors reviewed, edited, and approved the final version of the manuscript.

